# Opening the Pig to Comparative Neuroimaging: A Common Space Approach Contextualizes the Pig and Human Structural Connectome

**DOI:** 10.1101/2020.10.13.337436

**Authors:** R. Austin Benn, Rogier B. Mars, Ting Xu, Jason R. Yee, Luis Rodríguez-Esparragoza, Paula Montesinos, J.P Manzano-Patron, Gonzalo Lopez-Martin, Valentin Fuster, Javier Sanchez-Gonzalez, Eugene P. Duff, Borja Ibañez

**Affiliations:** Centro Nacional de Investigaciones Cardiovasculares (CNIC), Madrid, Spain; Université de Paris, CNRS, Integrative Neuroscience and Cognition Center, 75006 Paris, France; Wellcome Centre for Integrative Neuroimaging, FMRIB, Nuffield Department of Clinical Neurosciences, John Radcliffe Hospital, University of Oxford, Oxford, UK; Donders Institute for Brain, Cognition and Behaviour, Radboud University Nijmegen, Nijmegen, The Netherlands; Child Mind Institute, New York NY, USA; University of Veterinary Medicine, Vienna, Austria; Department of Neurology, Hospital Universitario Fundación Jimenez Diaz, Madrid, Spain; Department of Neurology, Stroke Unit, Hospital Universitari Germans Trias i Pujol, Badalona, Spain; Philips Healthcare, Madrid, Spain; Sir Peter Mansfield Imaging Centre, Mental Health and Clinical Neurosciences, School of Medicine, University of Nottingham, Nottingham, United Kingdom; Icahn School of Medicine at Mount Sinai, New York NY, USA; Department of Paediatrics, University of Oxford, Oxford, UK; IIS-Fundación Jiménez Díaz University Hospital, Madrid, Spain; CIBERCV, Madrid, Spain

## Abstract

Neuroimaging’s capability to quickly and rapidly phenotype the cortical organization of the whole brain brings with it the possibility to extend our understanding of cortical organization across the mammalian lineage. However, neuroimaging has thus far generally limited itself to a small number of species, with most animal studies being performed in either rodents or Non-Human Primates. Here we perform a first pass characterization of an animal which has recently seen its stock rise in the neuroscience community with the development of new models of neurological disease; the domestic pig. Characterizing the structural connectome of the pig, we create a white matter atlas, and an anatomical template which we use to build a horizontal translation between the pig and human based on a connectivity blueprint approach. We find that conserved trends of structural connectivity across species enabled spatial prediction of regions of interest between the pig and human, showing the potential horizontal translations have as a tool to assess the translational validity of porcine models of neurological disease. Releasing the anatomical template, white matter atlas, and connectivity blueprints, we hope to ease and promote the acceptance of the pig as an alternative large-animal model by the neuroimaging community.

## Introduction

The vast body of work performed in well-established animal models like rodents and Non-Human primates (NHP) may limit the widespread adoption of lesser studied animal models in neuroscience. Mechanistic studies prefer rodents since they are easy to handle and relatively easy to modify genetically, but large animals can better model certain aspects of human disease given their similar body size, organ shape, lifespan, and metabolism. These advantages are not exclusive to NHPs, and other large animal models, including pigs, dogs, and cats, all share a gyrencephalic brain, which can be studied using human imaging equipment. Years of tracer injections, imaging, and invasive recordings have provided a significant head start toward understanding the cortical organization of NHP’s. Recently interest has been growing in the use of the domestic pig (sus scrofa) as an alternative to NHP models^1^. Due to its popularity in biomedical research settings, the pig is already a widely available laboratory animal, and a new bevy of disease models have recently been developed using either genetic modification to create pigs with neurodegenerative diseases, including Parkinson’s, Alzheimer’s, and schizophrenia ^1,3,4^, or more traditional models of traumatic brain injury^5^, and stroke^6^. However, the tools and analysis workflows that make NHPs the premiere large animal model in neuroimaging today, are not present in the pig, limiting insight how cortical organization is altered in these recently developed models. To this end we create an open source package of utilities for neuroimaging in the pig (https://github.com/neurabenn/pig_connectivity_bp_preprint). The package includes a volumetric and surface based anatomical template space, a Diffusion-Weighted Imaging (DWI) derived white matter atlas, and a cross-species connectopic mapping between the pig and human cerebral cortex.

The aim of building this package is to enable the translation of results between the pig and human using multimodal MRI to create a “common space” of cross-species comparison^7^. The idea of a common space is well established in human neuroimaging pipelines which use a reference template brain such as the Montreal Neurological Institute (MNI152)^8^ to spatially normalize images acquired across multiple modalities for group-level statistics and comparisons. These “vertical” within-species translations permit insight into cortical organization at the population level across distinct modalities of magnetic resonance imaging^7^. While templates have been constructed for the pig neuroimaging community^9–11^, differences in breed, animal size, and age led us to build the Porcine Neurological Imaging Space (PNI50), a volumetric and surface based template space representing the average brain of 50 Large White pigs.

Extending the common space approach, Mars et al. propose creating “horizontal” translations^7^, where shared features of cortical organization define a common axis for cross-species comparison. One way these concepts have been implemented in non-human primates is through a “connectivity blueprint”^12^, which uses structural connectivity derived from Diffusion Weighted Imaging (DWI), to create a cross-species common space based on the cortical projections of white matter (WM) tracts present in both species^12^. Applying this framework to the pig, we first characterize the structural connectivity of 6 pigs using DWI and probabilistic tractography, creating a white matter atlas (Figure 1B,3). Tracts identified include the previously reported structures of the corticospinal tract^13–15^ limbic fibers^16^, forceps minor^17^, cerebellar penduncles^17^, the anterior thalamic radiation^18,19^, and association fibers including the uncinate, inferior longitudinal, and inferior fronto-occipital Fascicules^20^. Each white matter tract’s projection to the cortical surface is then used within the connectivity blueprint framework to attain cross-species translation between the primate and ungulate lineage (Figure 1B).

**Figure 1:**
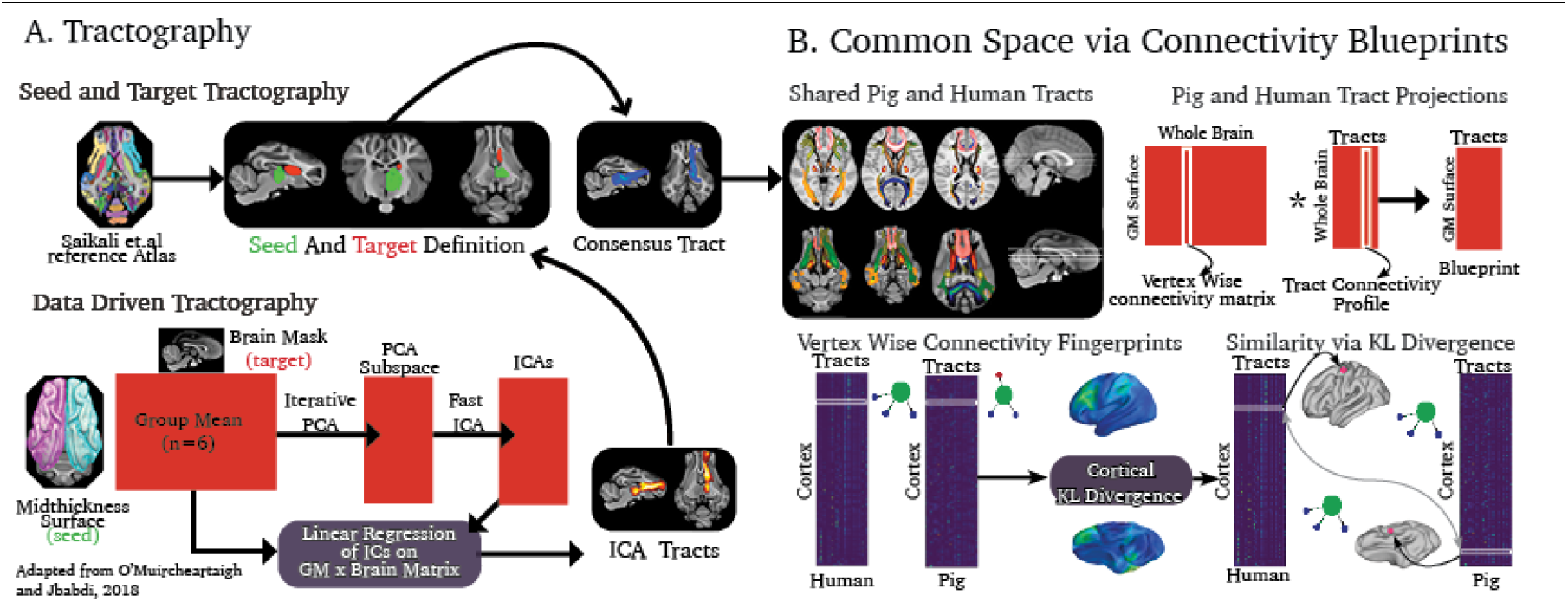
A). Data-driven tractography uses a gray matter surface seeding to a low-resolution volumetric target mask. PCA is then run on the resulting tractography matrix, followed by ICA in the PC subspace, and linear regression returns the tracts to their volumetric space. Tractography protocols were then defined using information gained from the resulting ICAs, and masks were drawn in the PNI50 space using the Saikali et al. atlas. All tracts, their protocols, and structures are described in the supplementary material. B). Identifying common tracts between the pig and human, we calculated how each tract connected to each vertex in the mid-thickness cortical surface. We then created a connectivity blueprint whereby the connectivity profile of each vertex to all the common tracts was stored in each row, and the tract cortical projection each column. The pig blueprint was used with a human blueprint with proposed common tracts, and a KL divergence similarity matrix was calculated to identify regions with the highest similarity across species.

A species’ connectivity blueprint consists of a collection of the cortical projections of its white matter tracts. Each vertex on the cortical surface is defined in terms of the tracts which reach it, i.e., its “connectivity profile,” and the overall collection of connectivity profiles for each vertex on the cortical surface is the connectivity blueprint. The connectivity blueprint is built through two separate tractographies (Figure 1B). The first is seeded from a cortical surface mesh and stores the streamlines propagated to a low-resolution volumetric brain mask in a *“surface x brain”* connectivity matrix. The second tractography targets the same volumetric brain mask but is seeded from tracts in the pig WM atlas to create a “tracts x brain” connectivity matrix (Figure 1B). The *surface x brain* matrix is multiplied by the *tract x brain* matrix to yield a connectivity blueprint (Figure 1B).

The connectivity blueprint of each species shifts the coordinates of the cortical surface mesh from representing a spatial to a connectopic position, defining each vertex by the tracts reaching it. The reframing of these coordinates is justified by the assumption that the mammalian connectome is organized to reduce wiring cost^21^, and that it does so through the principle of homophily^22^, or “like connects to like.” The mapping of each vertex’s position in the connectopic space permits the vertices in the pig and human to be matched by shared structural connectivity via the calculation of a similarity metric, in this case Kullback-Liebler divergence (KL). Calculating the KL divergence for every vertex between two species’ connectivity blueprints yields a cross-species similarity matrix that allows us to identify vertices with shared or divergent connectivity profiles, effectively mapping the pig and human cortices to one another (Figure 1B). Taking the minimum KL divergence across each axis of the similarity matrix, we can find where in the pig or human cortex structural connectivity has been conserved (Low KL) or where connectivity patterns have diverged (high KL) (Figure 1B). Finding where connectivity converges and diverges may then be used to build a translational interpretation of experimental results derived in pig models of neurological disease ^23^.

Until now the connectivity blueprint has been used within the primate lineage^12,24,25^. Through testing its limits we determine it can overcome 80 million years^26^ of divergent cortical evolution. This creates a translational framework whereby the pig and human can be contextualized in a common space such that the pig’s neurological disease models might find renewed relevance in their translational applicability to neuroscience.

## Results

### The Porcine Neurological Imaging Space (PNI50)

A factor contributing to the inertia in the adoption of lesser-used species such as pigs is that tools such as anatomical templates which permit group inference and vertical translation in human and NHP neuroimaging are often made in house, and seldom shared with the wider public. The Porcine Neurological Imaging Space (PNI50) is a standard reference space made from 50 anatomical T1-weighted images of male large white pigs weighing between 25-45 KG (figure 2A). Despite the large weight range in the pigs used to create the anatomical template, we found that spatial correlation using randomized sub-templates of smaller groups of pigs found limited anatomical variation, and that template stability could be achieved with as few as 5 pigs (Figure 2B). Spatial normalization to a volumetric template creates a space where traditional neuroimaging workflows can be used to identify features of cortical organization at a group level which might be used to create a horizontal axis for cross-species translation.

**Figure 2:**
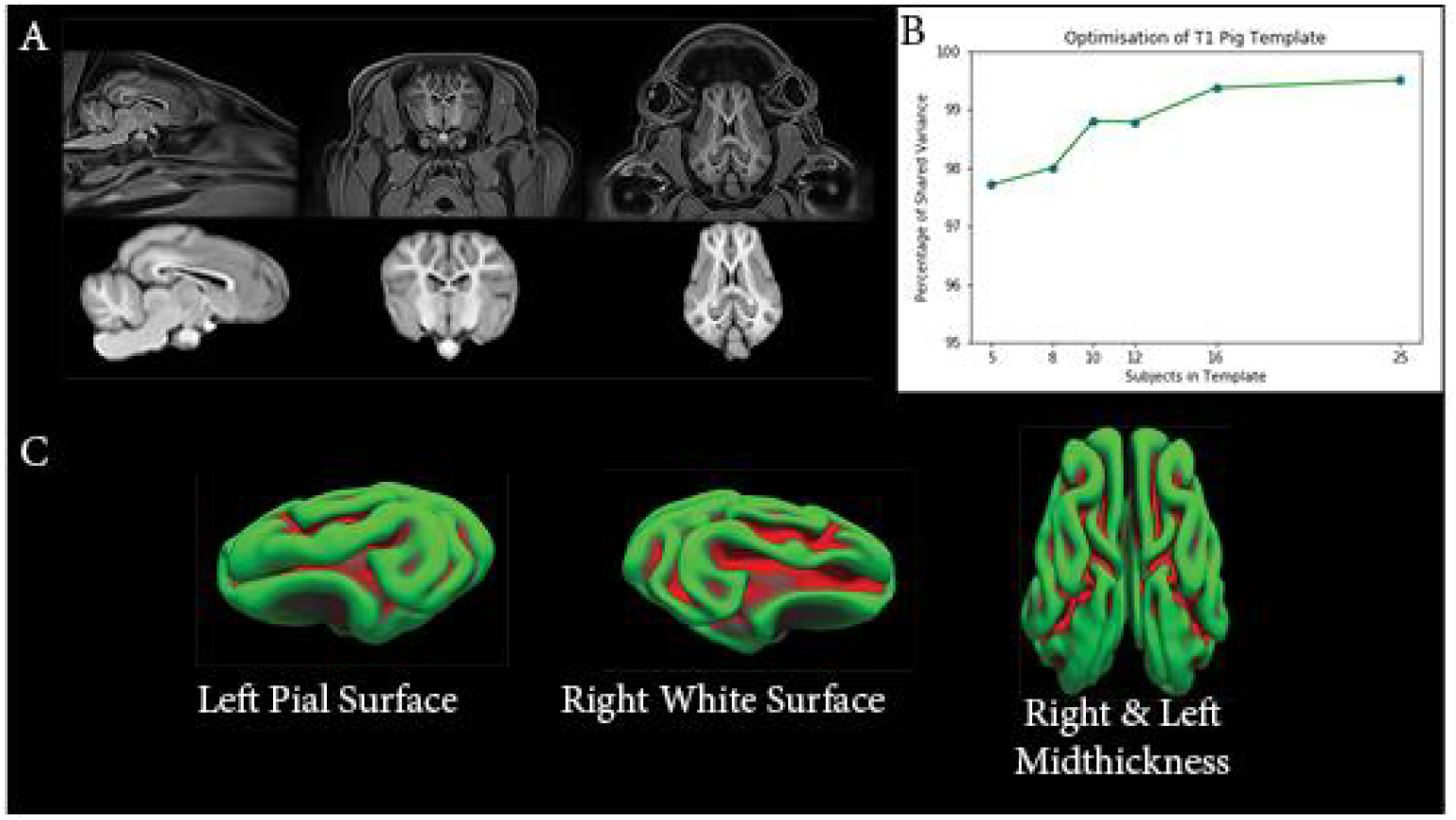
A). The whole-head and brain extracted PNI50 volumetric templates. B). Template optimization study shows the number of pigs needed to reach template stability. C). The PNI50 average surface space

However, volumetric templates have limited viability when creating a horizontal axis, as brain size, and the shape of each cortex inherently lead to a distinct dimensionality for each species. One way to overcome this is through cortical surface models, which can be decimated on a sphere to yield a common dimensionality for cross-species translation. To this end, we created an average surface template in PNI50 pace analogous to Freesurfer’s FsAverage (Figure 2C)^27^. With the average surface space, the remaining prerequisite to adapt the connectivity blueprint to the pig pertains to the identification of its white matter tracts.

### A Pig White Matter Atlas

Previous studies of the pig’s white matter have identified thalamic radiations, commissural, limbic, and association fibers^13,16,17,19,20^. Combining the information provided from prior work, and data-driven tractography^28^ we aimed to create a minimally biased set of tractography protocols compatible with FSL’s XTRACT^24^ package so they can be readily used by other groups working on diffusion MRI in the pig (Supplementary Material). As our primary goal is to understand how these tracts can be used to create connectopic mapping for cross-species translation via a connectivity blueprint, we describe the thalamic radiations, commissural, limbic, and association fibers (Figure 3), their tractography protocols, and the data-driven components which guide the tractography protocol’s definition in the supplementary material. Spatial descriptions of our results use the Saikali et al. gray matter atlas^29^ as a reference; however, we acknowledge that despite similar naming conventions, some labels may not indicate homology to structures in the human brain.

**Figure 3.**
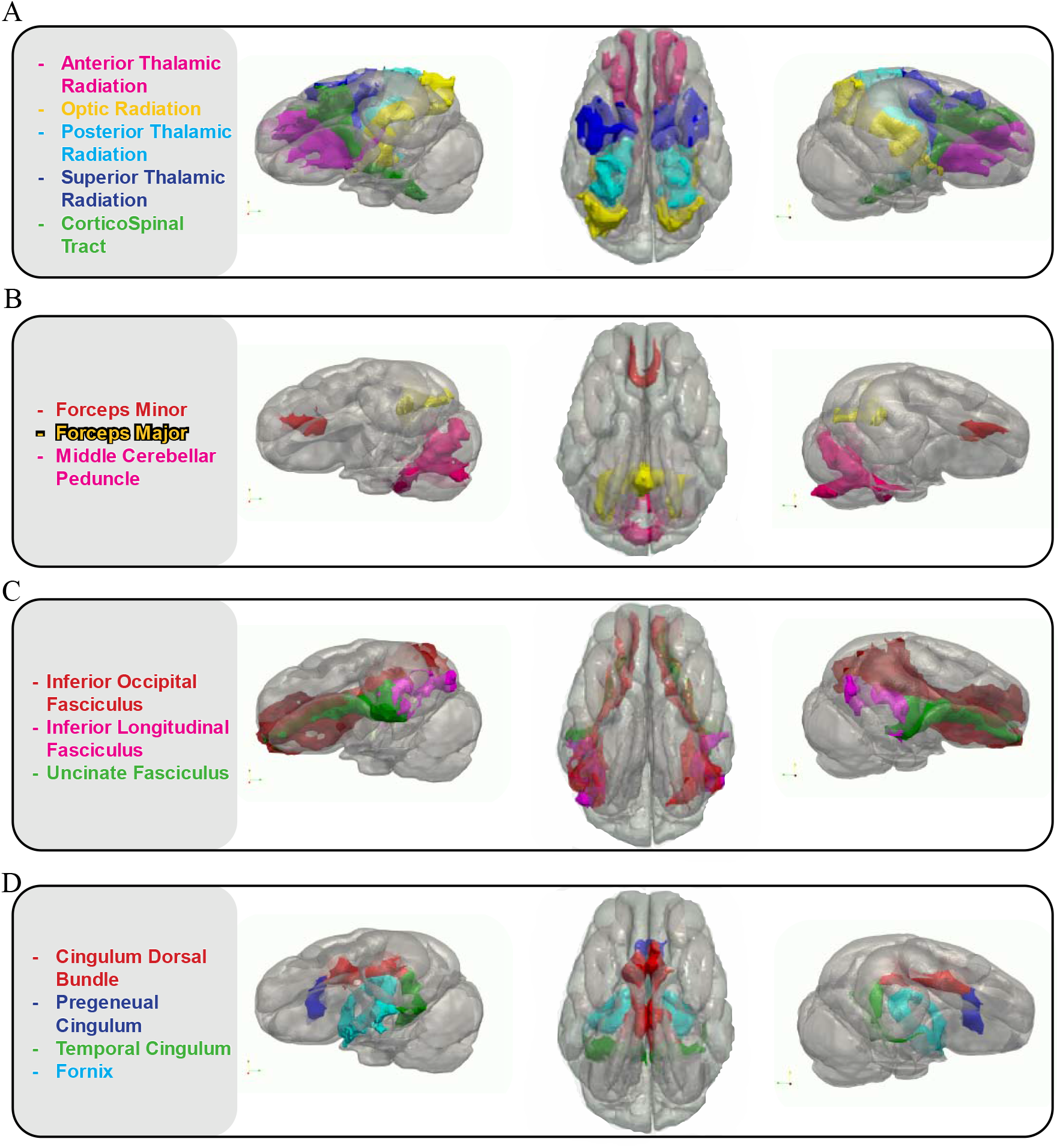
A). The Projection Tracts of the pig: the Anterior (ATR), Occipital (OR), Posterior (PTR), and Superior (STR) thalamic radiations and Corticospinal tract (CST). B). The Commissural and Cross-Hemispheric Tracts of the pig: the Forceps Major (FMA) and Minor (FMI), and the Middle Cerebellar Peduncle (MCP) reconstructed in 3D. C). The Association tracts of the pig: the Inferiorfrontal Occipital Fasciculus (IFOF), the Inferior longitudinal Fasciculus (ILF), and the Uncinate Fasciculus (UNC) reconstructed in 3D. D). The Limbic tracts of the pig: the Cingulum Dorsal Bundle (CBD), The Pregenual Cingulum (CBP), the Temporal Cingulum (CBT), and the Fornix (Fx) reconstructed in 3D

### Connectivity Blueprints

The connectivity blueprint can be thought of as a collection of white matter tract projections to the cortical surface. Each row of the connectivity blueprint contains the probability that a tract reaches a given vertex in the cortical surface mesh of each species. These connectivity profiles in each row form a diagnostic “connectivity fingerprint” defining each vertex through the tracts it connects to instead of spatial coordinates, creating a common connectivity space for cross-species comparison (Figure 1B). Given two connectivity blueprints, the distance between two given vertices across species can be calculated through Kullback-Liebler divergence (KL). Doing so for the whole cortex yields a similarity matrix mapping pig and human vertices within the joint pig-to-human structural connectivity space. This similarity matrix provides a basis not only for cross-species translation, but also gives a global perspective of where structural connectivity is convergent (low KL) or divergent (high KL).

### Connectivity blueprints spatially predict neural correlates between the pig and human

Twenty-seven tracts determined to have a similar course, start, and termination (supplementary material) were included in “global” connectivity blueprints of the pig and human. The connectivity blueprint of each species included the tract projections of the projection, commissural, association, and limbic fibers (Figure 1B, Figure 3) of each species (supplementary material). As an initial test to determine the viability of cross-species connectopic mapping between the pig and human, we used the KL divergence matrix calculated between the pig and human connectivity blueprint and used inverse-distance weighted interpolation to predict regions of interest defined in the Harvard/Oxford MNI152 atlas^30^ onto the pig cortex (Figure 4). Using masks from the frontal pole, occipital pole, and precentral gyrus, our spatial prediction of these regions in the pig aligns with the Saikali et al. atlas^29^ labels corresponding to the dorsolateral prefrontal cortex, V1 of the visual cortex, and the primary somatosensory cortex (Figure 4). Applying the inverse translation of the Saikali et al. labels, we find a reasonable alignment is obtained with the ROIs taken from the Harvard/Oxford atlas (Figure 5). Two-way predictions across species demonstrate the connectivity blueprint’s ability to create a horizontal translation between the pig and human cortex, a critical first step in expanding the pig’s role in neuroscience. Of course, we do not claim that pig and human frontal pole are functional homologs, as it is known that parts of this part of the human brain are a unique expansion in that lineage^31^, but rather that the connectivity blueprint framework and its derived horizontal translations shift the interpretation of homology to a nonbinary continuum which we for the first time extend to a large gyrencephalic animal model outside the primate lineage.

**Figure 4:**
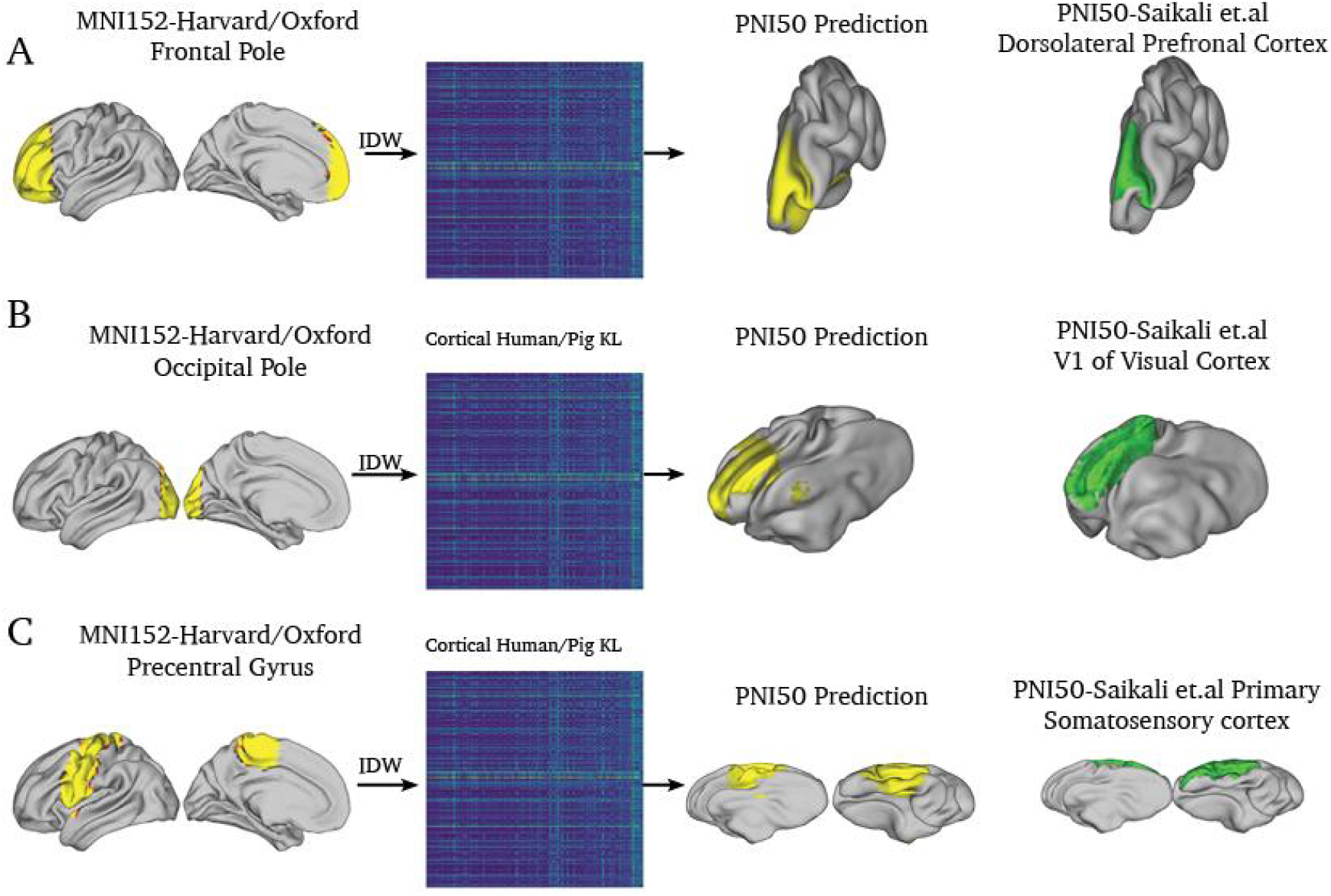
Spatial predictions of human regions of interest onto the porcine cortex using the KL similarity and Inverse Distance Weighted Interpolation (IDW)matrix calculated with the global connectivity blueprints The frontal pole, as defined in the Harvard/Oxford atlas, interpolated and predicted onto the pig (yellow) and the Saikali et al. DLPFC (green) in PNI50 space. The occipital pole in the Harvard/Oxford atlas interpolated and predicted onto the pig (yellow) and the ground truth V1 mask of Saikali et al. atlas in PNI50 space. The precentral gyrus, as defined in the Harvard/Oxford atlas, interpolated and predicted onto the pig (yellow) and the ground truth primary somatosensory cortex of the Saikali et al. atlas in PNI50 space.

**Figure 5:**
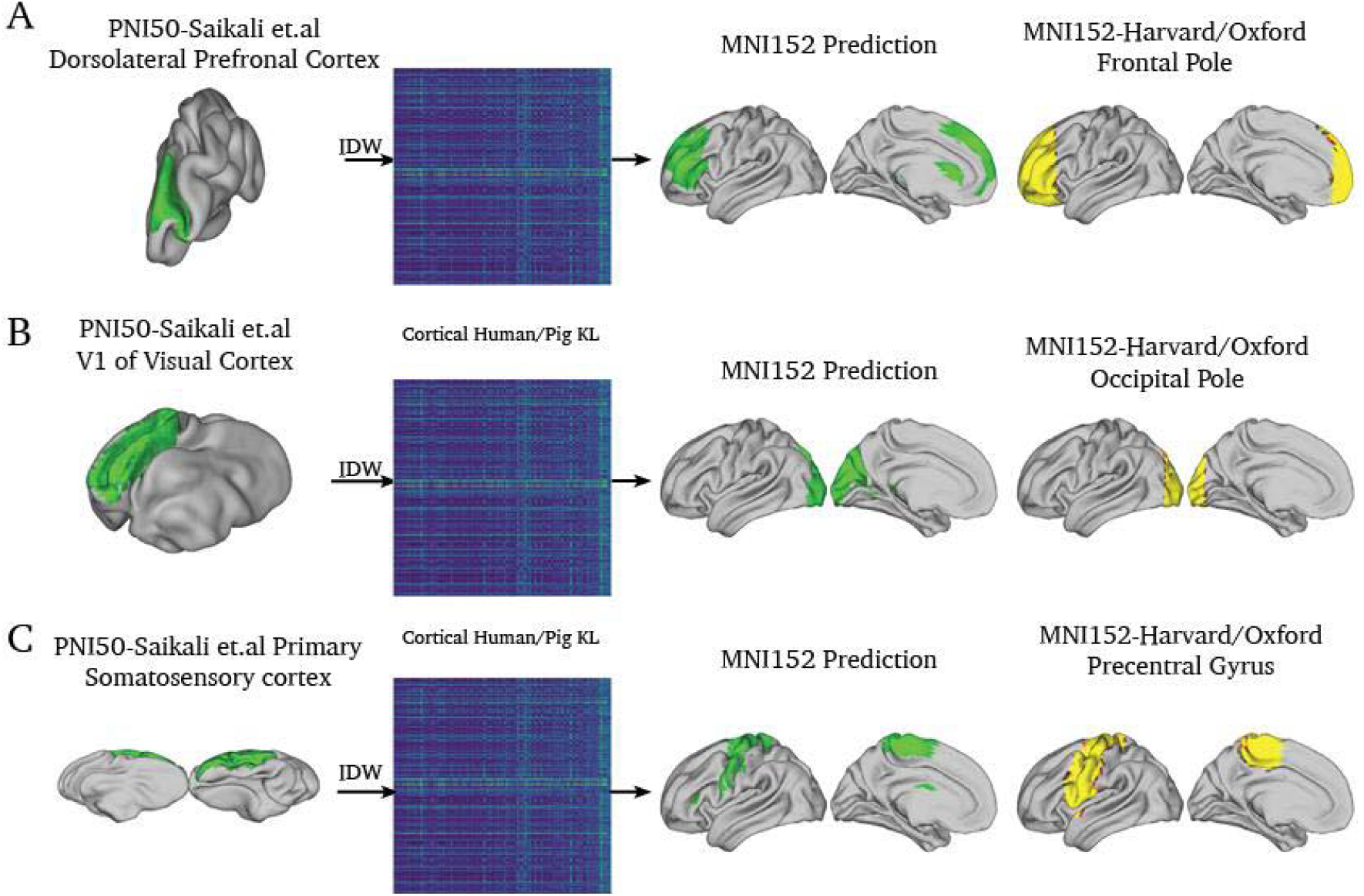
The inverse of figure 4, predictions of pig regions of interest onto the human cortex using the KL similarity matrix and Inverse Distance Weighted Interpolation (IDW) calculated with the global connectivity blueprints A). The Saikali et al. DLPFC (green) in PNI50 space interpolated and predicted onto the human surface (green) and the frontal pole as defined in the Harvard/Oxford atlas (yellow). B). The Saikali et al. V1 of the visual cortex (green) in PNI50 space interpolated and predicted onto the human surface (green) and the occipital pole as defined in the Harvard/Oxford atlas (yellow). C). The Saikali et al. Primary Somatosensory Cortex (green) in PNI50 space interpolated and predicted onto the human surface (green) and the precentral gyrus as defined in the Harvard/Oxford atlas (yellow).

### Identifying patterns of conserved connectivity based on tract projections to the pig and human cerebral cortices

Having identified tracts associated with major fiber bundles in the pig, we use the connectivity blueprint to ascertain the level to which a given group of tracts contributes to the conservation of connectivity across species. This was done with a leave-one-out analysis where a tract group (projection, commissural, association, and limbic fibers) is removed from the “global blueprint” containing all 27 tracts from our WM atlas. Visualizing the projections of each tract, we observe the contribution these morphological differences play in changing the minimum KL divergence across blueprints (Figure 6,7,8,9). This permits us to describe the spatial changes in the minimum KL divergence changes between the global blueprint, and the leave-one-out blueprints, quantifying the effect a tract group plays in the global min-KL via spatial cross-correlation (Figure 6,7,8,9 B). In the following section, we discuss the effect subtraction of a given tract group plays on the distribution of the minimum KL divergence between the pig and human, and explain it through the characterization of each tract’s projections to the cerebral cortex.

**Figure 6:**
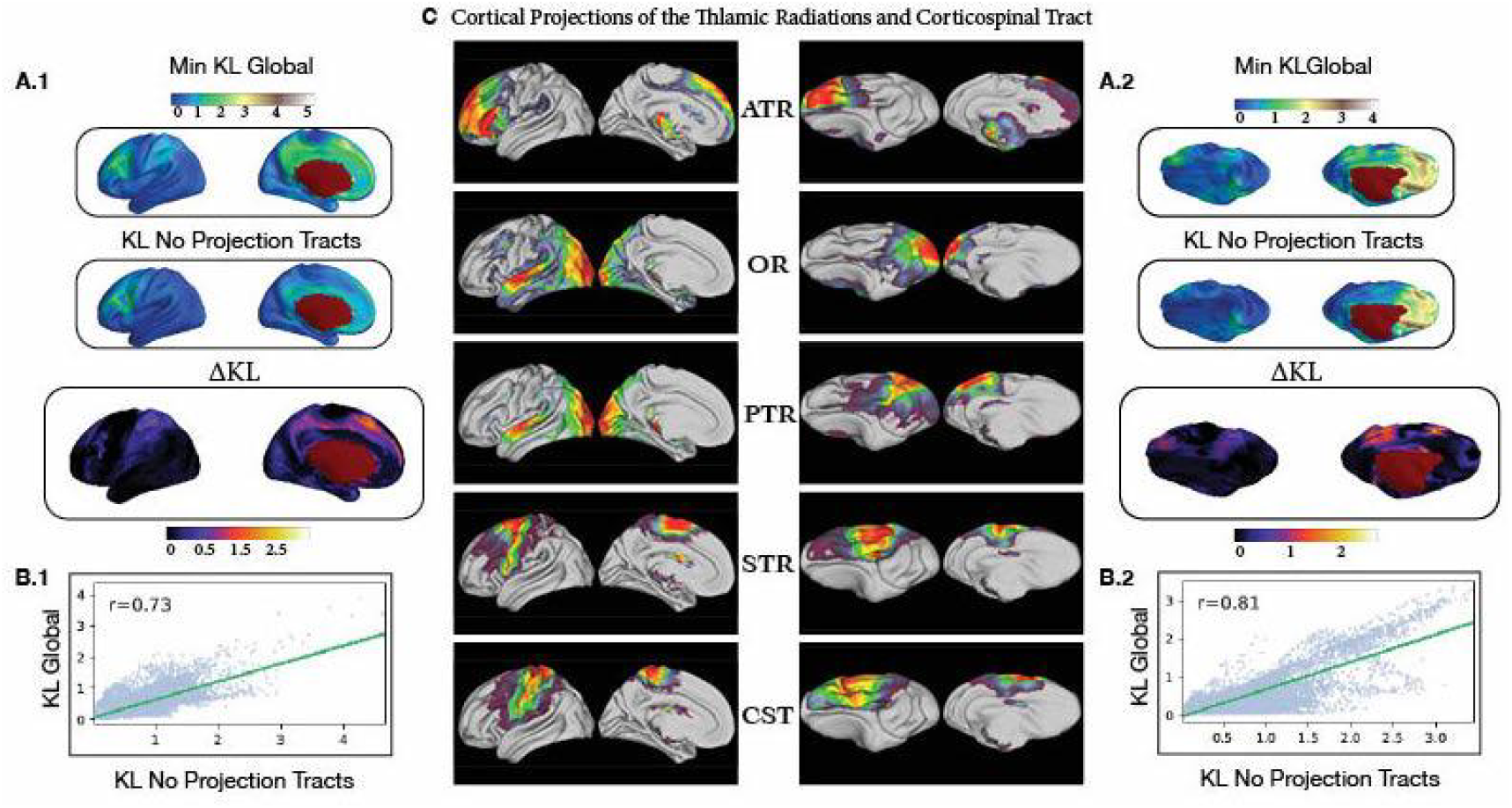
A.1). The projection tracts are removed from the global blueprint and the difference in minimum KL divergence corresponds to areas in the medial frontal cortex and the precuneus. A.2). The KL difference corresponds to the frontal division of the coronal sulcus where the frontal and somatosensory regions divide in the pig B.1) The projection tracts in the human blueprint significantly drive the overall KL divergence of the full blueprint as shown by a low spatial cross correlation of r=0.73. B.2) A significant portion of the KL divergence can be attributed to the presence of the projection tracts (r=0.81). C). The cortical tract projections of the Anterior (ATR), Occipital (OR), Posterior (PTR), Superior (STR), and Corticospinal tract (CST).

**Figure 7:**
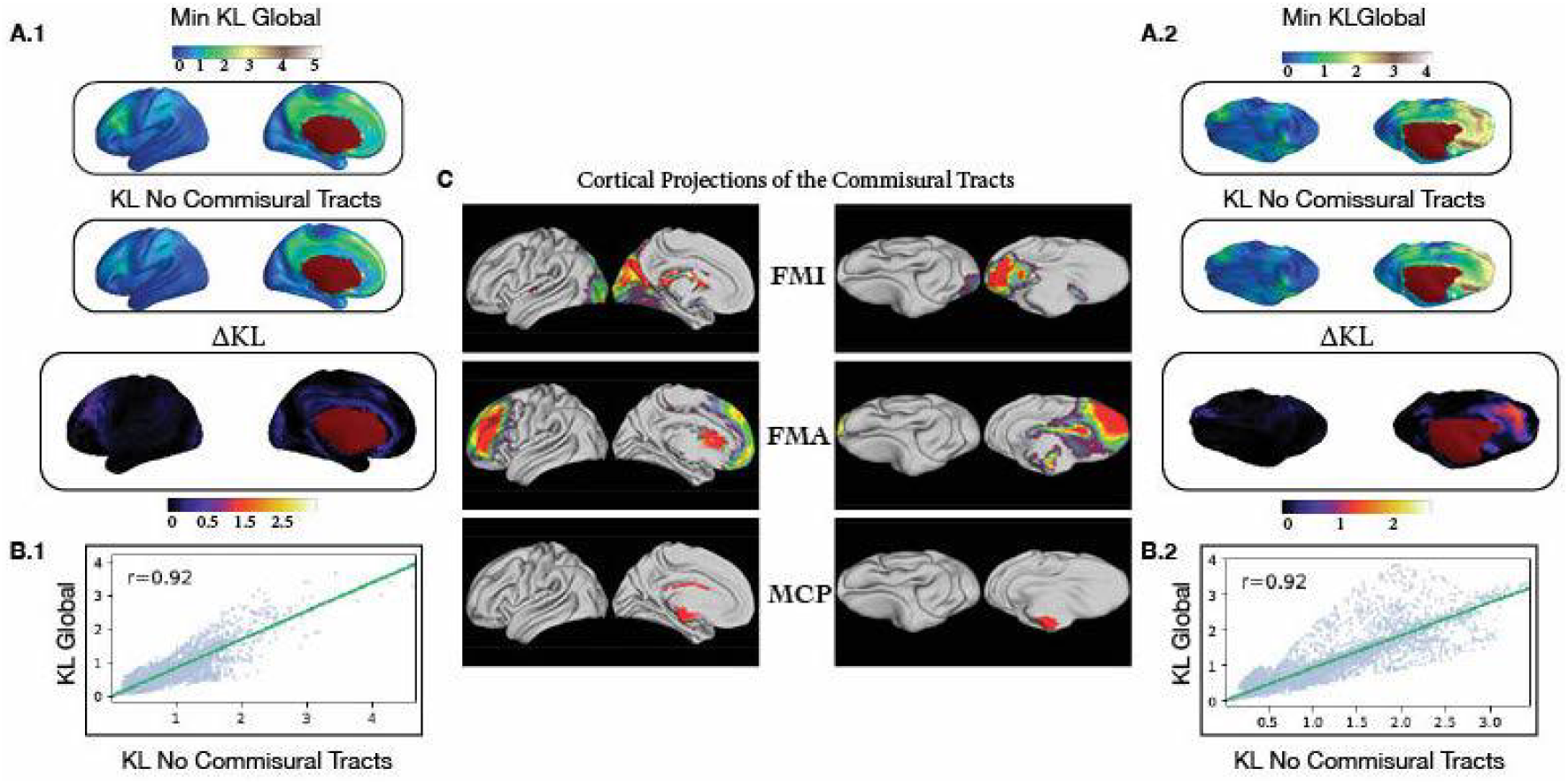
A.1) Mild changes in KL divergence are found in the region of the lateral projections of the Forceps Minor A.2) Changes in the KL divergence are shown in the medial territory of the FMI and Forceps Major B.1 & B.2). The impact of the commissural tracts is minimal at r=0.92 implying that these tracts are highly conserved between the pig and human and minimally alter the overall KL divergence between both species. C). The cortical tract projections of the Forceps Minor (FMI), Forceps Major (FMA), and Middle Cerebellar Peduncle (MCP).

**Figure 8:**
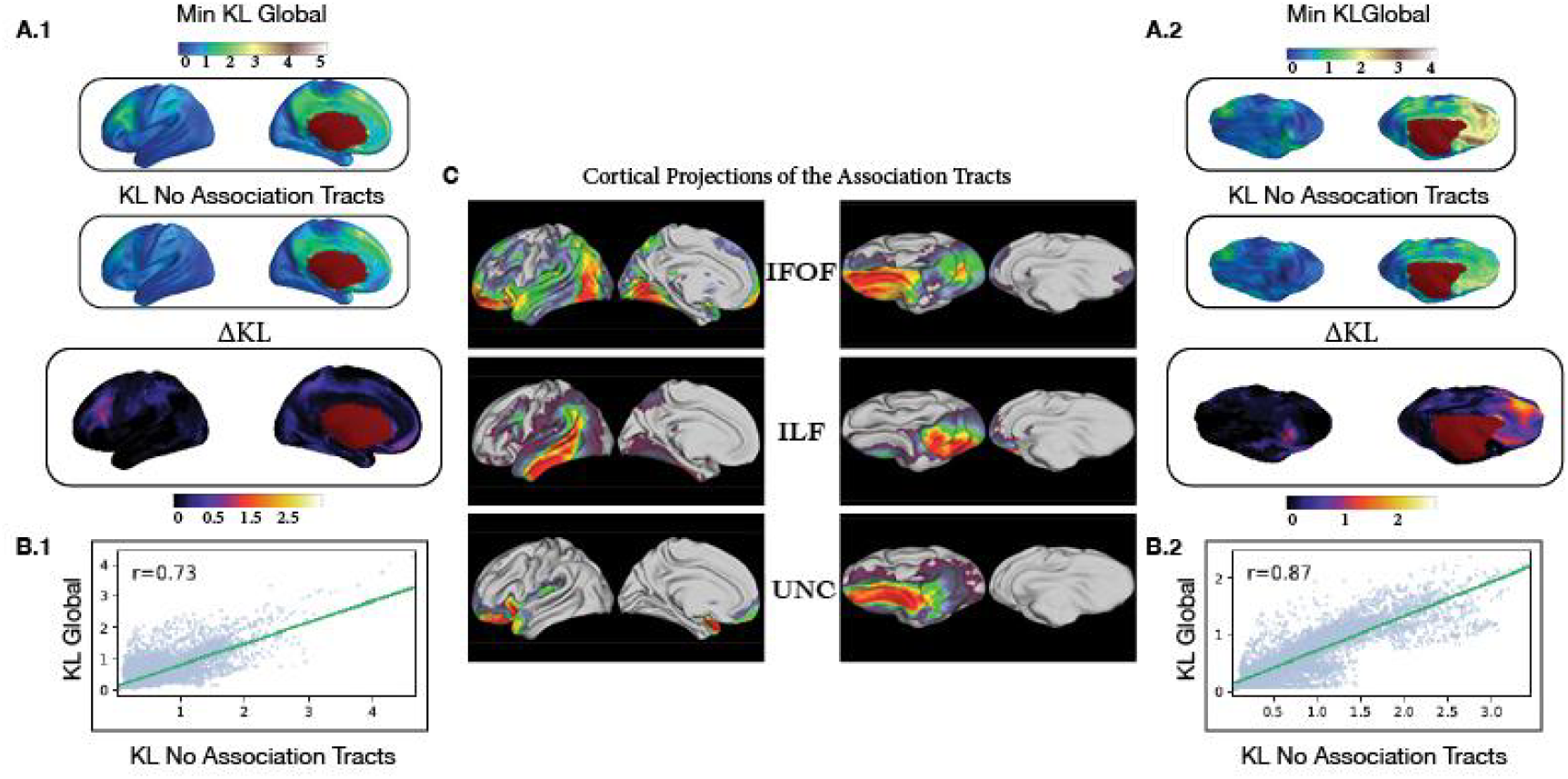
A.1) KL is observed to change in the medial frontal gyrus implicating the association tracts as responsible for the KL divergence values of the global blueprint A.2). The pig min KL changes in the superior frontal cortex along the medial surface, as well as the inferior temporal gyrus B.1). The association tracts lower the spatial correlation to r=0.73 showing they play a considerable role in driving the dissimilarity measured in the human cortex. B.2). The association tracts play a smaller role in driving the full blueprint KL divergence than the projection tracts (r=0.87) suggesting they may not be well conserved as their effect on the KL divergence of the full blueprint differs substantially for both species. C). The projection of the Inferior Fronto-occipital fasciculus (IFOF), Inferior Longitudinal Fasciculus, and Uncinate Fasciculus

**Figure 9.**
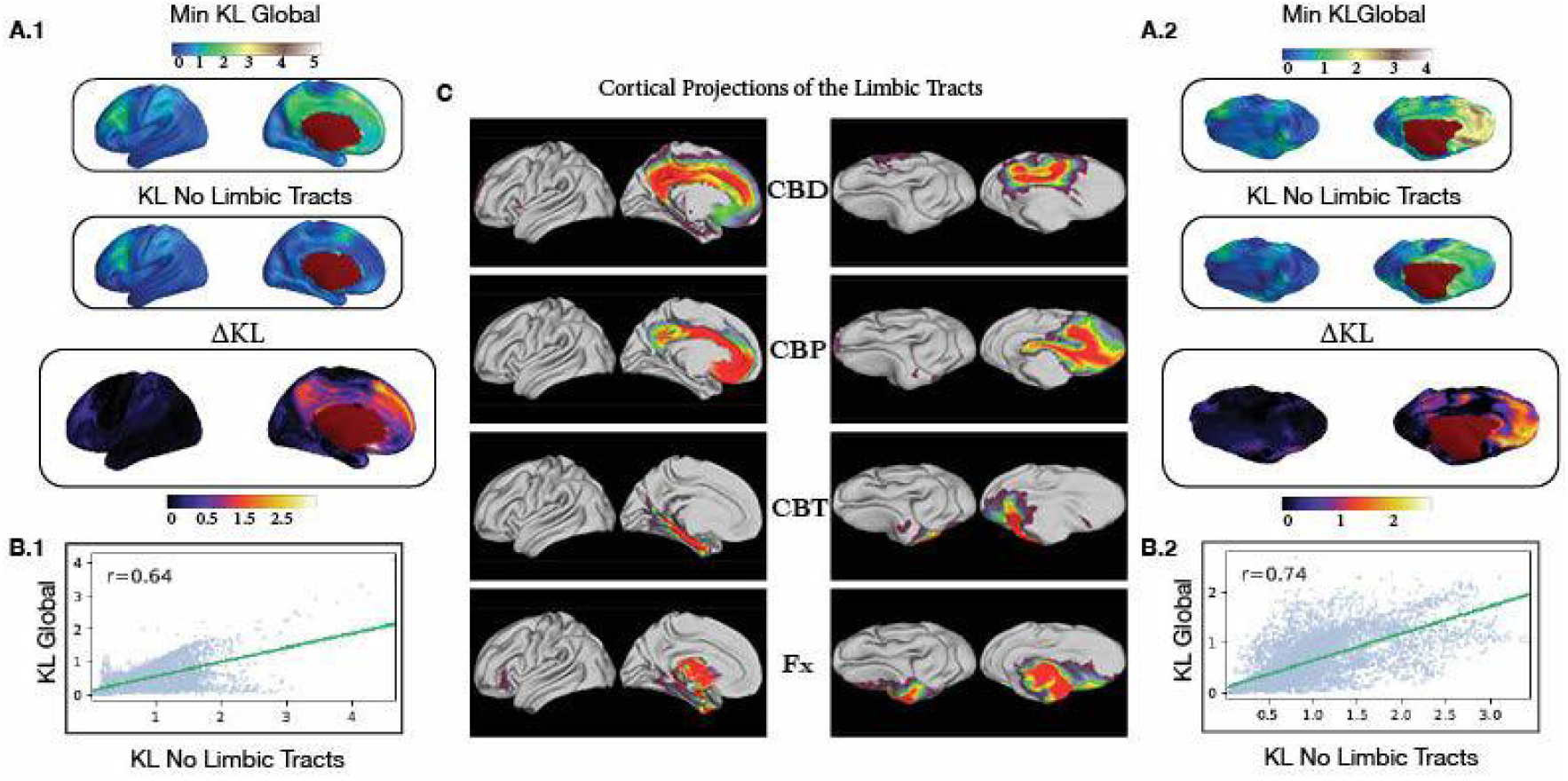
A.1) The KL divergence increases substantially along the body of the cingulum but not the fornix. The limbic tracts show a significant role in the divergence of the medial frontal cortex as well as in the precuneus A.2) The change in KL peaks in the frontal cortex of the pig as well as in the precuneus, but unlike the human does not outline a central body of the cingulum along the anterior-posterior axis B.1). The limbic tracts have the lowest spatial correlation (r=0.64) with the min-KL of the global blueprint B.2). The limbic tracts are the primary drivers of the KL divergence in the global min KL of the pig with a minimal spatial correlation (r=0.74). C). The cortical tract projections of the Cingulum Dorsal Bundle (CBD), the Pregenual Cingulum (CBP), Temporal Cingulum (CBT), and Fornix (Fx)

### The Projection Tracts

The lowest similarity between species in the projection tracts as indicated by the greatest change in KL divergence, corresponds to the tract projections of the ATR and CST (Figure 6A). The lateral division of the prefrontal and somatosensory cortices in the pig leads to partial innervation of the pig’s anterior somatosensory cortex by the ATR, a feature not present in the human. The cortical projections of the pig CST extend into the anterior frontal lobe overlapping with the projections of the ATR, whereas in the human, they remain within the pre and post-central gyrus (Figure 8A). The overlap between the CST and ATR of the pig causes an increase of unique connectivity fingerprints in the pig’s frontal lobe where the local minimum of KL-divergence peaks (Figure 6A). Increased KL-divergence is further observed in regions associated with the PTR and OR, given their increased presence in the temporal lobe of the human(Figure 6C). The overall role of the projection fibers in driving KL divergence is considerable as shown by the decrease in spatial correlation between the full blueprint and leave-one-out group (Figure 6B).

### The Commissural Tracts

The global KL divergence is mildly influenced in the regions innervated by the FMA and FMI (Figure 7A,C). The cortical projections of the FMA are conserved as they enter the medial occipital lobe in both species (Figure 7C). Similarly, the MCP is conserved through a lack of projections to the cortex in both species (Figure 7C). The FMI increases KL divergence in the lateral prefrontal cortex of the human and the medial-inferior frontal lobe of the pig (Figure 7A). However, the FMI’s cortical projection does not cross the lateral barrier into the somatosensory cortex, making it spatially conserved within the prefrontal cortex (Figure 7C). The overall impact of the commissural tracts as compared to the projection or other tract groups seems to be minimal as shown by its high spatial correlation with the global blueprint (Figure 7B).

### The Association Tracts

The min-KL divergence is primarily impacted in the prefrontal cortex and anterolateral somatosensory cortex (Figure 8A). Conversely, the change in KL divergence in the human cortical surface suggests that KL divergence in the Angular/Supramarginal Gyri and lateral prefrontal cortex is impacted by their absence (Figure 8A, 8C). If we assume the pig’s tract coursing from the occipital to the frontal cortex to be the IFOF, we observe that changes in the KL divergence are still associated with each species’ distinct IFOF projection patterns (Figure 8C). The pig’s IFOF projects to the cortical surface in the anterior prefrontal cortex and occipital lobe but lacks the temporoparietal projections found in the human IFOF (Figure 8C). Furthermore, the human IFOF innervates a far greater portion of the cortex, increasing the probability that any given point of the human cortex connects to the IFOF compared to the pig (Figure S5).

The UNC charts a course similar to that of the human but shows increased KL divergence in the temporal pole and inferior frontal cortex. The tract’s cortical projections remain close to the tract starting in the inferior temporal gyrus and ending in the inferior frontal lobe (Figure 8C), leading to what appears to be an elongated morphology as compared to that of the human (Figure 3D,8C). The pig’s lack of anterolateral expansion in the temporal lobe causes the pig ILF projection to run horizontally from the inferior to superior temporal gyrus, likely contributing to the changes observed in the min KL divergence in the pig’s inferior temporal lobe (Figure 8A). Conversely, where significant anterolateral expansion of the temporal lobe has occurred in the human as compared to the pig, the ILF’s cortical projection runs diagonally (Figure 8C).

The IFOF, ILF, and UNC share structural characteristics in both species. However, the reduction of spatial correlation between the global connectivity blueprint min KL sand the leave-one-out group shows these tracts introduce a greater proportion of non-mutual connectivity fingerprints between species than either the commissural or projection fibers. Notably, the gap in spatial correlation with the global KL divergence is more pronounced in the human cortex (Figure 8B), and can likely be attributed to the association tracts’ extensive cortical projections in the human brain as compared to those of the pig (Figure 8C, S5).

### The Limbic Tracts

Of the four white matter tract groups profiled here, the subtraction of the limbic fibers causes the greatest reduction of spatial correlation with the mink KL of the global blueprint in both species. While the fornix appears to have similar cortical projections across species, change in KL divergence is obvious in the medial frontal lobe and cingulum; territory belonging to the divisions of the cingulum bundle (Figure 9A, 9C). The pig does not appear to possess the continuity associated with the human cingulum, as made evident through each species’ cortical projections. Specifically, we note the pig CBD arches upwards, connecting the precuneus and somatosensory/premotor area complex instead of extending anteriorly into the territory of the CBP as in the human (Figure 9C). Without the continuum’s presence in the pig, the proportion of shared connectivity fingerprints decreases, causing the low spatial correlation between the global minimum KL divergence and the leave-one-out group(Figure 9B).

## Discussion

Understanding the pig’s structural connectome and contextualizing it with the human cortex using the connectivity blueprint approach provides an impetus to overcome the present inertia of its widespread adoption as an animal model in neuroscience. The pig has long been considered one of the premiere large-animal models for translational research in corporeal pathology as convergent evolution has led to a striking similarity in the morphology and function between the internal organs of the pig and human^32–36^. However, translation between the pig and human in neuroscience is a unique challenge, as cortical morphology has evolved on divergent paths to fill each species’ unique ecological niche^7^. To let other groups in neuroscience adapt the pig into their existing workflows as a gyrencephalic out-group in phylogenetic studies, or simply enabling spatial prediction of corresponding cortical regions between the pig and human, we release a package permitting the alignment, construction, and application of cross-species spatial prediction (https://github.com/neurabenn/pig_connectivity_bp_preprint) between the pig and human (Figure 4,5). This package includes the PNI50 average volumetric and surface-based templates, the white matter atlas and its XTRACT^24^ compatible tractography protocols (Supplementary Material), and the connectivity blueprints redefining the pig and human coordinates to a common connectopic space.

The global connectivity blueprint is comprised of 27 tracts and was used to spatially predict the coordinates of atlas defined regions of interest across the pig and human, establishing a horizontal axis of common connectivity between the pig and human (Figure 1B, 4,5). Combined with the leave-one-out analysis, each of these predictions provides us valuable information as to the differences in cortical morphology, and how common, and divergent connectivity might inform how we can interpret results derived from the pig’s cerebral cortex in an experimental setting.

For example, cross-species prediction and lower KL divergence values were found across the visual cortex of both species. This was unexpected, as although primary cortex is considered to be well conserved across the mammalian lineage^37^, the pig’s visual system contains a panoramic field extending 310° degrees on each side as opposed to the primate’s forward-looking visual system which prioritizes binocular vision and depth perception^37,38^. However, despite these stark differences we were able to predict V1 of the pig to the human and of the human to the pig accurately (Figure 4,5), in a region where the KL divergence was relatively stable (Figure 6) despite innervation by the IFOF, ILF, PTR, OR, and FMA (Figure 6C, 8C). One possibility is that we did not include a Ventral Occipital Fasciculus (VOF) in the pig WM atlas. Future work may include the identification of a pig VOF, and its validity within the connectivity blueprint could be assessed as to whether it contributes to convergent, or divergent connectivity with the occipital pole across species. Still, the high degree to which structural connectivity was conserved in different visual systems, and across the leave-one-out analyses suggests structural connectivity alone may not be sufficient to identify divergent functional specialization. Thus, to fully adapt the pig as a validated model of neurological disease, future work will have to characterize the functional organization of the pig cortex (Figure 6,7,8,9).

The spatial prediction of the somatosensory and the prefrontal cortex was also successful, despite distinct morphology in the separation of the pig and human somatosensory and frontal lobes. Whereas the human sensory-frontal division runs anterior-posterior across the central sulcus, the pig’s sensory-frontal division is orthogonal to that of the human, splitting laterally across the coronal sulcus^29^. Cross-species spatial predictions reflected this difference suggesting the connectivity blueprint provides a means of cross-species mapping in accordance with the cytoarchitecturally derived boundaries of each species (Figure 5,6)^29^. That these boundaries were respected was surprising as although the STR and CST innervate the somatosensory and motor cortex in both species, the CST and STR extend into the pig’s frontal lobe, and the pig’s ATR further diverges from human connectivity patterns, entering the anterior somatosensory cortex (Figure 7C). Our ATR is in accordance with previous tracer studies of the pig’s^18^, but its prior use as a defining feature of the prefrontal cortex^19^ suggests again that further study of the pig’s functional organization may provide insight as to whether the somatosensory region in the frontal lobe corresponds to sensory, or association cortex.

Spatial prediction of the frontal lobe across species was relatively successful, and we might attribute this to relatively conserved tract projections from the ATR, IFOF, FMI, and UNC (Figure 6,8). However, our leave one out analysis shows that unique connectivity fingerprints are incurred with the inclusion of the association tracts in the lateral prefrontal cortex (Figure 8B), likely due to the distinct projections of the human IFOF to the parietal and temporal lobe (Figure 8C). The Uncinate Fasciculus of the pig is flatter and possesses a distinct angular orientation as compared to the human (Figure 8A,C), and while this difference appears to minimally affect KL divergence across the species, we posit the pig UNC’s unique tract structure may be due to minimal anterolateral expansion of the temporal lobe as compared to the human cortex (Figure 3C). Within the temporal lobe, we do find ILF and UNC structures, but also note that unlike the human, the optic and posterior thalamic radiations do not enter the temporal lobe (Figure 6C), and moreover, the pig IFOF does not maintain the presence of widespread temporal and parietal projections (Figure 8C). Given these differences in tract morphology, we feel it would be premature to claim the pig’s IFOF as homologous, but considering homology on a spectrum as implied by the measure of KL divergence, it is worth considering that a ventral longitudinal tract has been identified in at least 130 mammalian species^21^.

Divergence between the global KL divergence and our leave-one-out blueprints shows the association tracts drive a significant portion of the global KL divergence (Figure 8B), but that the limbic tracts most drive the difference between the pig and human connectivity blueprints (Figure 9B). This was surprising, as the tracts of the limbic system were reconstructed with data-driven methods^28^, and our mask protocols produced similar structures to the recent work of Bech et al.^16^ (Figure 3D). Most of the change in KL divergence as compared to the global blueprint can be attributed to the cingulum bundle, as unlike the human, the pig’s cingulum tracts do not form a continuum as evidenced through the discontinuous tract projections of the CBD, CBP, and CBT (Figure 9C). This change is a major deviation from the human structural connectome, but their partial derivation by data-driven methods (Figure 1A, Supplementary Material) and their correspondence to the work of Bech et al., and our ability to successfully perform spatial prediction across species (Figure 4,5) suggest they deserve their place in the connectivity blueprint.

## Limitations

The tracts included in the pig and human connectivity blueprints are generally considered symmetrical in the human brain. Without the Superior Longitudinal (SLF) and Arcuate Fasciculus in both blueprints, cortical symmetry of connectivity may be overestimated in both species (Figure S5). While we found data-driven components suggesting SLF-like structures and Pascalau et al. identify an SLF in a pig blunt dissection study^20^, we could not reconstruct the SLF with a hand-defined tractography protocol, leading us to leave it out of the final connectivity blueprint. Postmortem study at high resolution instead of the in-vivo 1.4 mm scans we performed might also provide a more precise delineation of the pig’s cingulum and detangle the overlapping cortical projections of the IFOF and UNC in the external capsule. Additionally, we note a significant discrepancy in data quality between our DWI data and the Human Connectome Project^39^ data used to generate human connectivity blueprint. However, the tracts derived align with those previously proposed in the literature^13,17,19,20^, suggesting that our resolution was sufficient for an initial study of the pig’s structural connectivity.

Although we have sought to match tracts included in the pig’s blueprint based on their similarity in their course and termination points, the labels we use here are merely suggestive of homology. Our approach uses suggested homology to investigate similarity and difference across brains, but it can also test explicit hypotheses of homology. For instance, Roumazeilles et al. showed that defining one branch of the human ILF as homologous to macaque ILF led to a greater overall divergence score than when defining another branch as homologous^40^. This suggests that only one of the human ILF branches is similar to that of the macaque. A similar approach can be employed to formally test homology of all proposed tracts here. However, it should be noted that the labeling of tracts with similar names is only the start of the investigation in the present approach, and indeed we can show that some tracts whose homology is more certain like the forceps major, and minor have a minimal impact on divergence scores.

## Conclusion

One promise of neuroimaging is that it allows for the rapid and comprehensive phenotyping of cortical organization. In comparative neuroanatomy these tools have provided insight into the evolution of cortical organization, but their scope has been limited to a few select species. We create a set of open-source tools that provide a new take on a gyrencephalic large animal model, the pig. Extending the *connectivity blueprint approach* beyond the primate lineage, we study the relation of cortical tract projections between the pig and human, finding that despite 80 Million Years of divergent cortical evolution, structural connectivity is sufficiently conserved in the sensory and association cortex to consider the pig as a model worthy of wider adoption in neuroimaging. Notably, the pig’s inclusion into the neuroimaging literature may yet provide us with a unique lens to identify traits unique to the primate cortex, while also extending the translational viability of findings derived in porcine models of neurological disease.

## Methods

The institutional animal review board approved all studies. Images were acquired in free-breathing pigs in the prone position on a Philips 3T Achieva scanner, with a Philips 32 channel Cardiac Coil (Philips Health Care, The Best Netherlands). All modalities had the same anatomical image sequence acquired, which we will describe here in the context of creating the Porcine Neurological Imaging Space.

### Animal Preparation for Diffusion-Weighted and Anatomical MRI

All images were acquired as part of baseline MRI studies from planned cardiology experiments in the lab. Anesthesia in these animals was induced with a cocktail of Ketamine (20 mg/kg), Midazolam (0.5 mg/kg), and Xylazine (0.2 mg/kg), and maintained via continuous intravenous infusion of ketamine (2 mg/kg/h), midazolam (0.2 mg/kg/h), and xylazine (0.2 mg/kg/h).

### Structural T1 Image Acquisition

Each pig studied in this thesis had a T1 weighted 3D Flash image acquired(TR 10 ms, TE 4.8 ms, Flip Angle(FA) 10°, FOV 210 mm, Matrix 264 × 238, 150 slices at 0.8 mm isotropic resolution).

### Construction of the Volumetric PNI50 template

A common space for spatial normalization and group analysis was constructed using the T1-weighted images of fifty male pigs weighing between 25-50Kg. Pigs were initially registered linearly using FSL’s FLIRT to a single subject using 12 degrees of freedom(dof)^41,42^. Following initial alignment, the antsMultivariateTemplate.sh script was run to create a “whole-head” anatomical template^43^. We next defined a brain mask in the “whole-head” template space that was warped to all 50 pigs and used for brain extraction. The ANTs template script then runs on only the brain extracted images.

### Template Optimization study

To determine the stability of volumetric templates in the pig, we performed a brief template optimization study where we created sub-templates ranging from 5 to 25 pigs. The fslcc tool was used to calculate the spatial correlation of each sub-template with the other sub-templates containing the same number of subjects. We report the mean value of each template’s within-group spatial correlation as a measure of template stability, as recently reported by Croxson et al. in the primate cortex^44^.

### The PNI50 average surface space

The preclinical surface processing pipeline *precon_all* (https://github.com/neurabenn/precon_all) run on the PNI50 volumetric template and the 50 pigs used in its construction. Precon_all is a pipeline created for the generation of cortical surface meshes in preclinical animal models which combines FSL^41^, Freesurfer^27^, ANTs^48^, and the human connectome workbench^45^. The pipeline can be run using a single T1-weighted image, and outputs a directory structure analogous to Freesurfer’s recon-all. Following the generation of white, gray, midthickness, and spherical surfaces, the surface of the PNI50 volumetric template were used as an initial template surface and each individual’s surfaces underwent iterative spherical registration. After registering the surfaces to one another, surfaces were averaged to create the PNI50 average surface space, a space analogous to Freeusurfer’s FSaverage.

### DWI Image Acquisition

Diffusion-weighted data was acquired with 64 encoding directions and a single B0 image (TR 13,500 ms, TE 100.5 ms, Flip Angle (FA) 10°, FOV 210 mm, Matrix 148 x, 150 slices, Slice Thickness 70, Slice Gap 0, Resolution 1.4 mm isotropic). Field map correction was done with a blip phase-encoded Image of the same geometry and acquisition parameters as the Diffusion-weighted Image.

### Preprocessing of DWI data

DWI images were first corrected for susceptibility-induced distortions with FSL topup and then corrected for movement and eddy current off-resonance effects in FSL eddy^46,47^. DWI images were rigidly registered to their anatomical images in FSL FLIRT with 6 dof^42^. The linear registration was concatenated to the nonlinear warp FSL FNIRT generated in the brain extraction process of precon_all^55^. The concatenated warp was then inverted, and the standard brain mask resampled into the native DWI space for brain extraction.

We then prepared the data for probabilistic tractography by running a modified version of the automated preprocessing pipeline autoPtx^48^. This included the generation of deterministic tensors and Fractional Anisotropy images using DTIfit and was followed by a probabilistic two-fiber tensor model estimation in BedpostX^48–50^.

### Data-Driven tractography

Using the average mid-thickness cortical surface of each hemisphere as a seed and a low resolution (1.4 × 1.4 mm) whole-brain target mask, we generated a matrix of streamlines passing from each vertex to every voxel in the brain. This was done through the probtrackx2, --omatrix2 option, and a step length specified to 0.35. After doing this for all 6 pigs, we performed iterative principal components analysis (iPCA) on the group average *surface x brain* matrix generated in the previous step. The results of iPCA were then fed into an Independent Components Analysis with a dimensionality set to 50 to identify exploratory tracts in the surface space. Linear regression then mapped the components on the surface space back into volumetric space, whereby they were saved and used to guide the tractography protocol definition. This was done with the moriarty and lookatmoriarty MATLAB scripts from the Mr Cat toolbox (www.neuroecologylab.org)^28,51^ (Figure 1A). We only identified 27 tracts in our pig WM atlas, despite setting the dimensionality of the ICA to 50 components. However, given the nature of ICA to split components, it was common to find ICAs which contained only part of what was likely the whole tract structure. On the other hand, it was also common for derived ICAs to contain multiple tracts, such as in the ILF and FMA. All 50 ICAs of the left and right hemisphere are available as part of data and code release(https://github.com/neurabenn/pig_connectivity_bp_preprint) for visual inspection and for guidance in defining future tractography protocols.

### Tractography with mask protocols

Using both the Saikali et al. Atlas and the data-driven components, we defined tract protocols in the PNI50 space, which could be used with either the AutoPtx and XTRACT packages^24,48^. These packages allow for tractography masks defined in a standard space to be warped to a subject’s individual space for tract reconstruction using Probtrackx, significantly streamlining the process of identifying tracts across multiple subjects. Using the default options of Probtrackx in autoPtx, we successfully reconstructed 27 tracts in 6 pigs. We then took each individual’s tract and transformed it back into the PNI50 space, where the average of all normalized streamlines was taken as the final group tract for the WM atlas.

### Constructing a Connectivity Blueprint for the pig

The connectivity blueprint is a *cortical surface* × *tract matrix* describing the connectivity fingerprint between each vertex of the grey/white matter surface and the tracts it connects to (Figure 1B). The connectivity blueprint is created by the multiplication of a *surface x brain matrix* and a tracts by *brain matrix*^12^. The group *surface x brain matrix* was generated in step one of the data-driven tractography. The *tract x brain* matrix is formed by a second tractography where each tract is used as a seed, and the same low-resolution mask from data-driven tractography is the target once again with the –omat2 option specified in Probtrackx. The mean of this output is taken and added to a matrix containing the mean tract to brain connections of the other tracts identified in the pig WM atlas to form a *tracts x brain matrix*. The connectivity blueprint for each hemisphere was then generated by multiplying the *tract* × *brain* and *surface* × *brain* matrices to produce a collection of the white matter tract projections to the cortical surface (figure 1B, 8)^12^.

### Human Connectivity Blueprints

Human connectivity blueprints and surfaces were obtained from Mars et al.’s original connectivity blueprint paper (https://git.fmrib.ox.ac.uk/rmars/comparing-connectivity-blueprints)^12^.

### Cross-Species Prediction of Regions of Human Interest

Using the global connectivity blueprint containing all 27, we calculated the KL similarity matrix for every cortical vertex between species. Regions of interest were projected to their respective surface. An inverse weighted distance interpolation was applied to each region using the similarity matrix and code modified Mars et al. (Figure 4,5)^12^.

### Identifying the tract Groups differentiating the pig and human

The pig connectivity blueprints contain 27 tracts, and the human blueprint had 39 tracts. All normalization and calculations of KL divergence were performed as in Mars et al.^12^. Using a python implementation of the code, we removed all tracts in the human blueprint not present in the pig. Having normalized the tracts in the blueprints, we calculated the KL divergence between the pig and human cortex. KL divergence measures the relative entropy between two probability distributions and quantifies the amount of information lost when using one distribution to estimate the other. Lower KL divergence means less information is lost, and thus greater similarity between both distributions. In this case, mapping the minimum KL divergence over the human and pig surface provides a visual representation of regions where the connectivity fingerprints between the pig and human are the most and least similar, where high KL signifies significant organizational changes between species.

Connectivity Blueprints consist of a collection of *connectivity fingerprints* of the whole cortex where the probability of each vertex connecting to each white matter tract is recorded. To better understand the contribution of each WM structure to the overall KL divergence, we performed a leave-one-out analysis whereby a single group of tracts was removed from the global blueprint. To visualize where on the cortex the minimum KL divergence changed we subtracted each leave-one-out blueprint minimum KL from the global minimum KL, creating a spatial map of where diverging connectivity fingerprints were present on the cerebral cortex. Furthermore we quantified the relation between the global blueprint and the leave-one-out groups via spatial cross-correlation which provides us with a quantitative measure of how each tract group contributes to the overall KL divergence. All calculations for the leave one-out KL divergence of tract groups are available in the data and code release (https://github.com/neurabenn/pig_connectivity_bp_preprint) as part of an interactive Jupyter notebook.

## Author contributions

RAB: contributed to Conceptualization, Methodology, Data Acquisition, Formal analysis, Visualization, Writing - original draft, Writing - review & editing; RBM: contributed to Conceptualization, Methodology, Formal analysis, Writing - review & editing; TX: contributed to Methodology, Analysis, Visualization, Writing - review & editing,; JY: contributed to Writing - review & editing LRE: contributed to Methodology, Data Acquisition, Formal analysis; PM: contributed to Methodology, Data Acquisition; JPM: contributed to Data Acquisition, Formal analysis, Writing - review & editing; GLM: contributed to Data Acquisition; VF: contributed to Writing -review & editing, JGS: contributed to Conceptualization, Data Acquisition, Visualization, Writing - review & editing; ED: contributed to Conceptualization, Methodology, and Writing - review & editing; BI: contributed to Conceptualization, Methodology, Data Acquisition, Writing - review & editing;

### Materials & Correspondence

Materials and correspondence can be directed to either R. Austin Benn or Borja Ibañez

