## Supplementary Material for "Opening the Pig to Comparative Neuroimaging: A Common Space Approach Contextualizes the Pig and Human Structural Connectome"

**Data-driven tractography Guides the definition of Tractography Protocols:**

Following the development of the PNI50, we describe the pig's structural connectome and identify 27 white matter tracts in the pig, which share a similar course and termination points white matter tracts in the human. To minimize prior bias in the construction of each tract protocol, we employ a data-driven tractography seeding from the PNI50 average mid-thickness surface to a volumetric low-resolution whole-brain target mask (1.4mm isotropic)(Behrens et al., 2007). The output of this tractography consists of a matrix containing the streamlines mapping each cortical vertex to every brain voxel(O’Muircheartaigh and Jbabdi, 2018). This matrix then underwent iterative Principal Components Analysis (iPCA) and Independent Components Analysis (ICA) (Figure 1A). Volumetric tractograms were then generated via linear regression of the ICA spatial maps back into volumetric space, where they were visually assessed as plausible tracts (Figure 1A)(O’Muircheartaigh and Jbabdi, 2018). Using the volumetric ICA tractograms as a guide, seed and target masks were hand-drawn in the PNI50 standard space (Figure 3-6). Each tractography protocol consists of a set of masks containing: the *seed* (tractography start point), *target* (waypoint, only streamlines that travel through here are retained), *exclusion* (areas prohibited to the streamline), and occasionally, *stop* (stops streamline propagation). All tractography protocols except those of the cross-hemispheric structures use the sagittal midline as an exclusion mask to prevent streamline propagation into the contralateral hemisphere. These tract protocols allow for reproducible tractography via compatibility with the automated tractography pipelines AutoPtx(De Groot et al., 2013) and XTRACT(Warrington et al., 2019). The repository presented here (<https://github.com/neurabenn/pig_connectivity_bp_preprint>) includes the 27 tractography protocols of the tracts used to build the pig's connectivity blueprint, as well as the corresponding data-driven ICAs for each tract(O’Muircheartaigh and Jbabdi, 2018; Saikali et al., 2010b).

**Projection Fibers:** Exploratory tractography found multiple components associated with the projection fibers, which we used to guide the definition of hand-drawn protocols; the whole thalamus was used as the seed region for all four thalamic radiations presented here. Based on principles of conserved cortical organization in the mammalian brain-plan(Krubitzer, 2007b) we expect the thalamocortical projections and corticospinal tract to serve as a base for comparison of conserved connectivity across species.

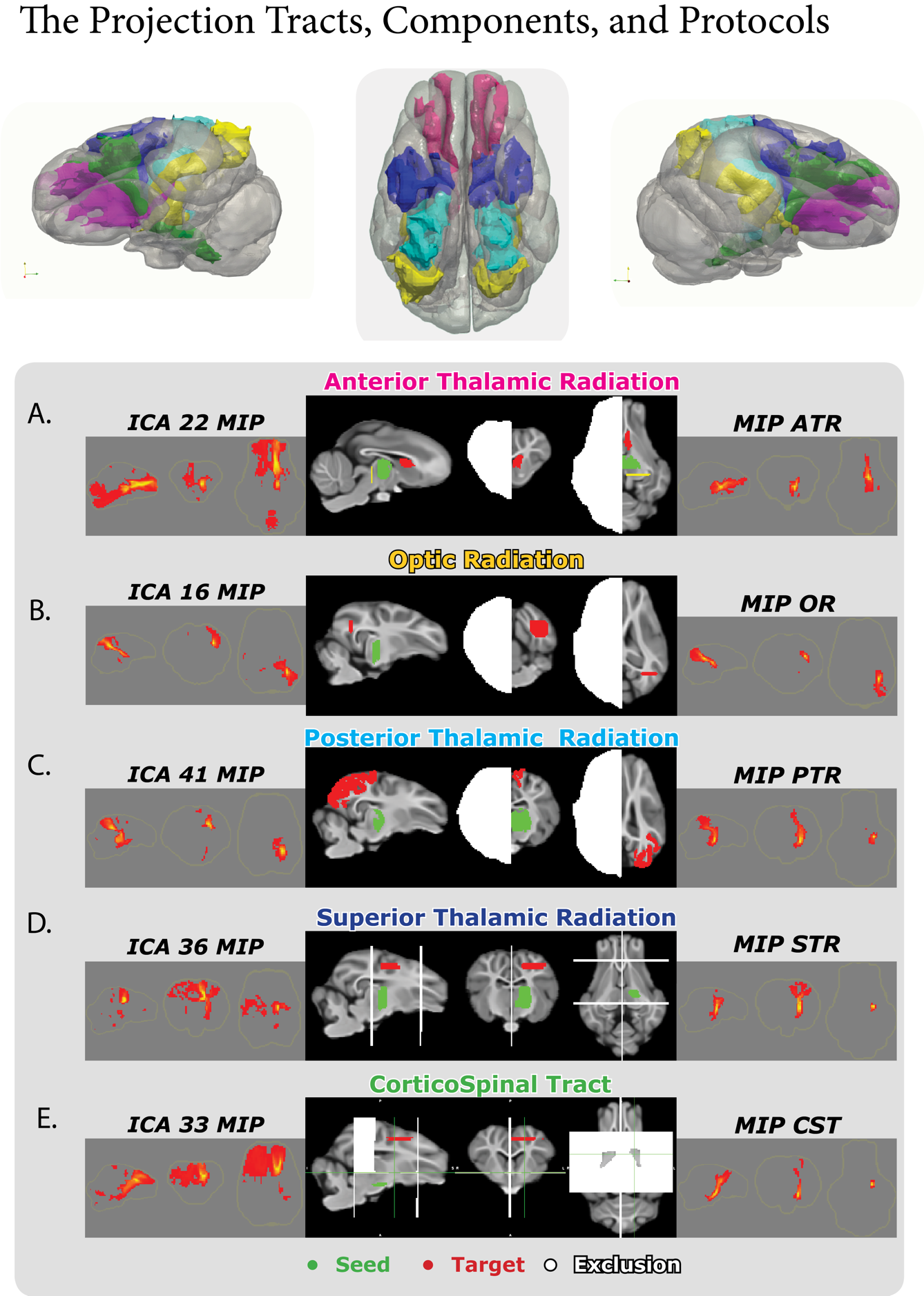

Figure S1: The Projection Tracts of the pig and human, including the Anterior

(ATR), Occipital (OR), Posterior (PTR), and Superior (STR) thalamic radiations and

Corticospinal tract (CST) are visualized as 3D reconstructions

A). The maximum intensity projection (MIP) of component L-22, the tractography protocol

used for the final reconstruction, and the MIP of the ATR reconstructed with the

mask protocols. B). The MIP of component L-16, the tractography protocol, and the

MIP of the OR. C). The MIP of component L-41, the tractography protocol, and the MIP

of the PTR. D). The MIP of component L-36 is associated with the STR, the tractography

protocol, and the MIP of the STR. E). The MIP of component L-33, the tractography

protocol, and the MIP of the CST.

*Anterior Thalamic Radiation (ATR):* Component 22 of the left hemisphere (L-22) and 45 in the right (R-45) contain a structure connecting the thalamus with the prefrontal cortex forming the ATR(Dyrby et al., 2007) (Figure 3). Using these components to guide the definition of the tractography protocol for the ATR, the target mask was defined as the caudate nucleus, and a coronal stop mask was drawn below the thalamus at the level of the posterior commissure (Figure 3A).

*Optic Radiation (OR):* The optic radiation OR connects the inferior visual lobe with the thalamus and is present in components L-16/41 and R-1/20. The target mask is a coronal slice in the inferior junction of V1 and V3 at the occipitotemporal junction (Figure 3B).

*Posterior Thalamic Radiation (PTR):* The PTR runs superior to the OR and connects the thalamus to a V1 target mask. The PTR was present in components L-13/25/39 and R-6/15(Figure 3C).

*Superior Thalamic Radiation (STR):* The Superior thalamic Radiation found in components L-17/36 and R-8/22/31 radiates from the thalamus to the primary and somatosensory association cortex Figure (7,8D). The target is an axial section superior to the seed region, where the Primary/Associative Somatosensory Cortex meet. Coronal slices anterior to the genu of the corpus callosum and at the posterior commissure are excluded (Figure 3D).

*Corticospinal Tract (CST):* The CST is partially found across components L-0/31/33/43 and R-0/23/42/19 (Figure 3E). The CST connects the brainstem and sensorimotor cortex, replicating the structure of Bech et al. (Bech et al., 2018b). The CST seed is drawn inferior to the thalamus, and the target is an axial slice of the sensorimotor cortex. The exclusion mask has two coronal slices at the posterior commissure, the genu of the corpus callosum, and a third axial slice inferior to the thalamus that does not include the internal capsule (Figure 3E).

**Commissural and cross-hemispheric Tracts:** The commissural fibers and the cerebellar peduncle were found in components that span both hemispheres (Figure 4). All three tracts used a reverse seeding strategy whereby the target and seed were flipped in separate reconstructions, and the final tract is presented as their average. All three protocols replicate the tracts previously identified by Zhong et al. (Zhong et al., 2016a).

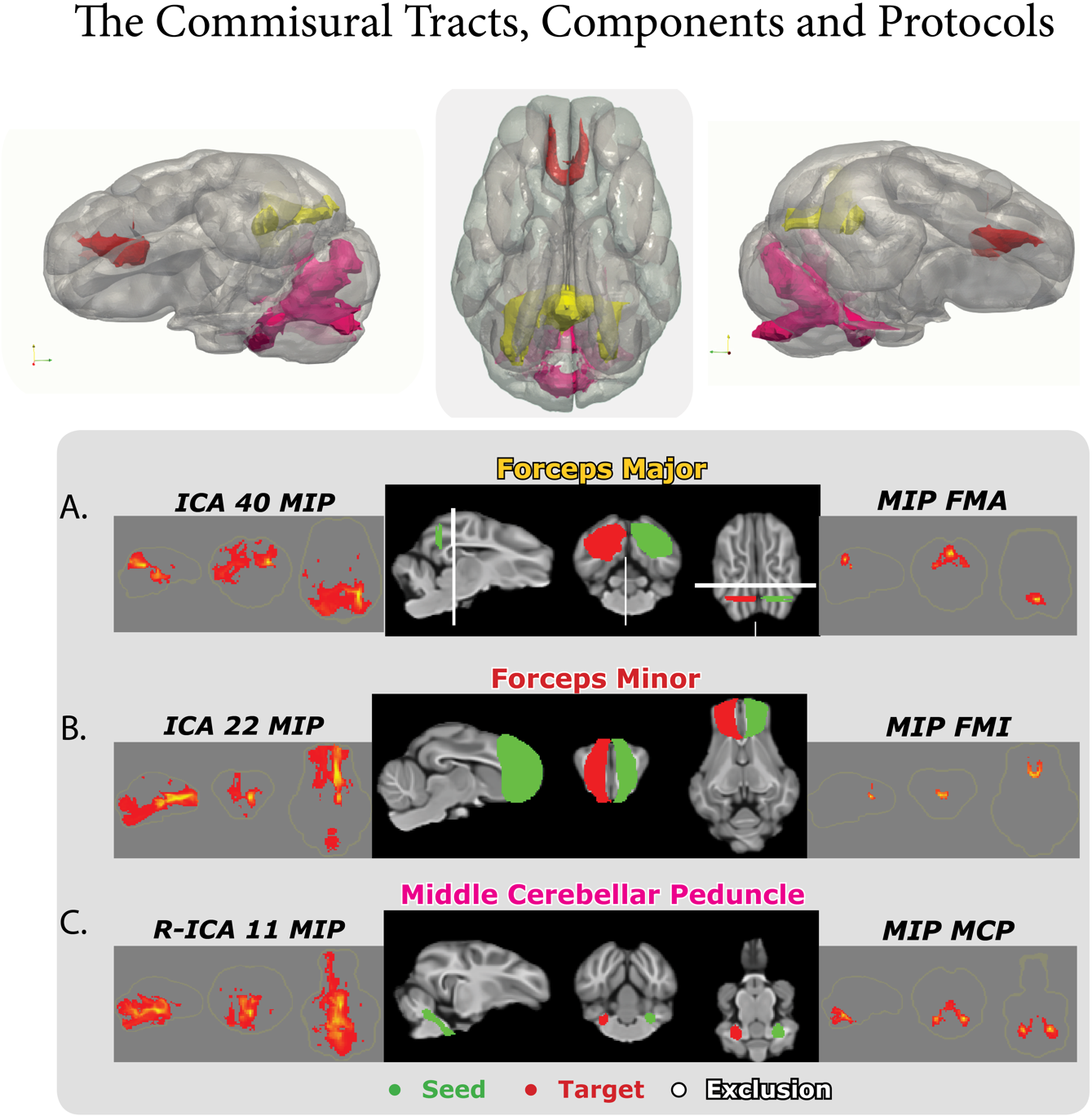

Figure S2: The Commissural and Cross-Hemispheric Tracts of the pig and human,

including the Forceps Major (FMA) and Minor (FMI), and the Middle Cerebellar Peduncle

(MCP) reconstructed in 3D.

A). The maximum intensity projection (MIP) of component L-40, the tractography protocol

used for the final reconstruction, and the MIP of the FMA reconstructed with the

mask protocols. B). The MIP of component L-22, the tractography protocol, and the

MIP of the FMI. C). The MIP of component R-11, the tractography protocol, and the

MIP of the MCP.

*Forceps Major (FMA):* The FMA (components L-40 and R-20) connects the left and right visual cortices passing through the splenium of the corpus callosum(Zhong et al., 2016a). The FMA was reverse seeded from V1 and V2 of each hemisphere, and a coronal section anterior to the splenium of the corpus callosum was excluded (Figure 4A).

*Forceps Minor (FMI):* The FMI connects the left and right prefrontal cortex, as shown in components L-4/22 and R-3/39 (Figure 4B). Coronal sections of the left and right dorsolateral prefrontal cortex are reverse-seed for the FMI, and there is no exclusion mask (Figure 4B).

*Middle Cerebellar Peduncle (MCP):* The MCP connects the left and right cerebellar peduncles and is found in components L-6/32 and R-42, which contain multiple tract structures(Zhong et al., 2016a). The MCP used the left cerebellar peduncle as a seed and the right as the target (Figure 4C).

**Association Fibers:**

Exploratory analysis with data-driven tractography identified three tracts with trajectories similar to the Inferior Fronto-Occipital Fasciculus, the Uncinate Fasciculus, and the Inferior Longitudinal Fasciculus of Pascalau et al.'s white matter dissection study (Figure 5)(Pascalau and Szabo, 2017). Components from data-driven tractography found structures reminiscent of the Superior Longitudinal Fasciculus (SLF) from Pascaleu et al.'s white matter dissection(Pascalau and Szabo, 2017). Our attempts to define a hand-drawn SLF tractography protocol were unsuccessful in replicating the SLF-like data-driven components, and we chose to leave it out of the pig white matter atlas for now.

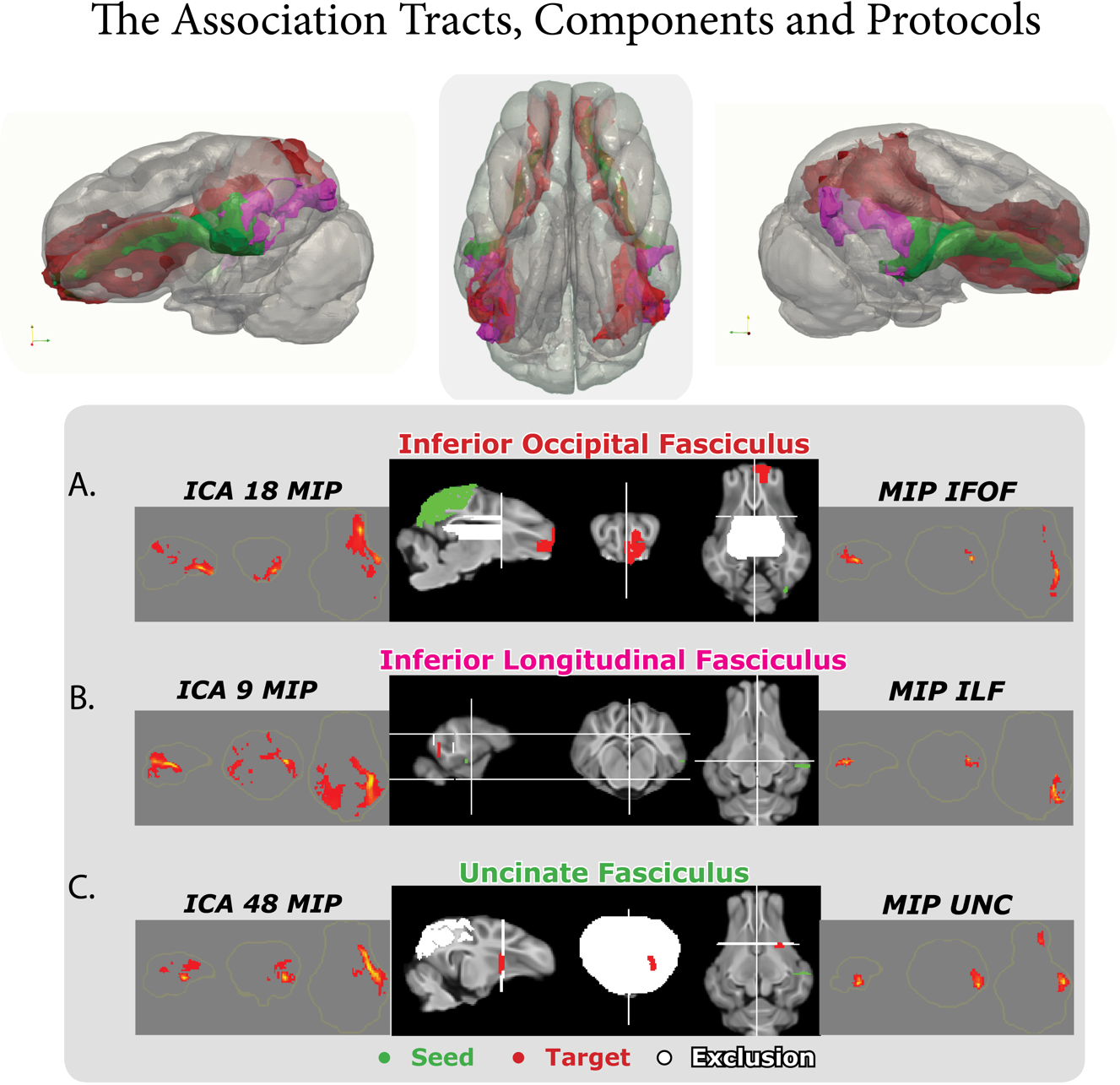

Figure S3: The Association tracts of the pig and human, including the Inferiorfrontal

Occipital Fasciculus (IFOF), the Inferior longitudinal Fasciculus (ILF), and the

Uncinate Fasciculus (UNC) reconstructed in 3D.

A). The maximum intensity projection (MIP) of component L-18, the tractography protocol

used for reconstruction of the IFOF, and the MIP of the IFOF reconstructed with

the mask protocols. B). The MIP of component L-9, the tractography protocol, and the

MIP of the ILF. C). The MIP of component L-48, the tractography protocol, and the MIP

of the UNC.

*Inferior Fronto-Occipital Fasciculus (IFOF):* Exploratory tractography found multiple independent components containing IFOF-like structures connecting the visual and frontal cortex. In the left hemisphere, these structures were present in components 34 and 18 (L-34/18) and component 21 (R-21) in the right hemisphere (Figure 5A). These components guided the IFOF seed placement using a visual cortex composite mask of V1, V2, and V3. The target mask was a coronal slice of the anterior prefrontal cortex, and the external capsule and the ventricles were excluded in the axial plane (Figure 5A).

*Inferior Longitudinal Fasciculus (ILF):* A tract reminiscent of the primate ILF is present as a substructure in components L-7/9 and R-15/20 (Figure 5B). The ILF connects the inferior temporal gyrus to the inferior occipital lobe. The seed mask is placed at the middle/inferior temporal gyrus, and the target included the superior temporal gyrus in an axial section at the level of the zona incerta. Coronal slices anterior to the seed and posterior to the target were excluded (Figure 5B).

*Uncinate Fasciculus (UNC):* The UNC connects the anterior region of the inferior temporal gyrus, arcs into the external capsule, and terminates in the Anterior Prefrontal Cortex. The UNC (L-48 and R-14/30) is seeded in the inferior temporal gyrus, where it targets the junction of the external capsule and putamen. The exclusion mask includes the visual cortex and the coronal plane around the target mask (Figure 5C).

**Limbic Tracts:** Data-driven tractography found components similar to fornix and cingulum, aligning with the recent work of Bech et al. (Bech et al., 2020b). As in primates, the cingulum bundle tractography protocols were defined in three parts: the temporal, dorsal, and pregenual bundles. ICA components of the data-driven tractography found three distinct structures of similar course and length to the cingulum proposed by Bech et al., suggesting a segmented reconstruction approach is necessary to reconstruct all three branches of the pig cingulum (Figure 6)(Bech et al., 2020b; Heilbronner and Haber, 2014). A notable difference between the pig's cingulum projections and those of the primate is that the pig cingulum does not project within itself to form a continuum(Figure 8)(Bubb et al., 2018). All protocols for the cingulum were reverse seeded. The fornix appears to be highly conserved across species as the primary hippocampal tract is derived from data-driven components.

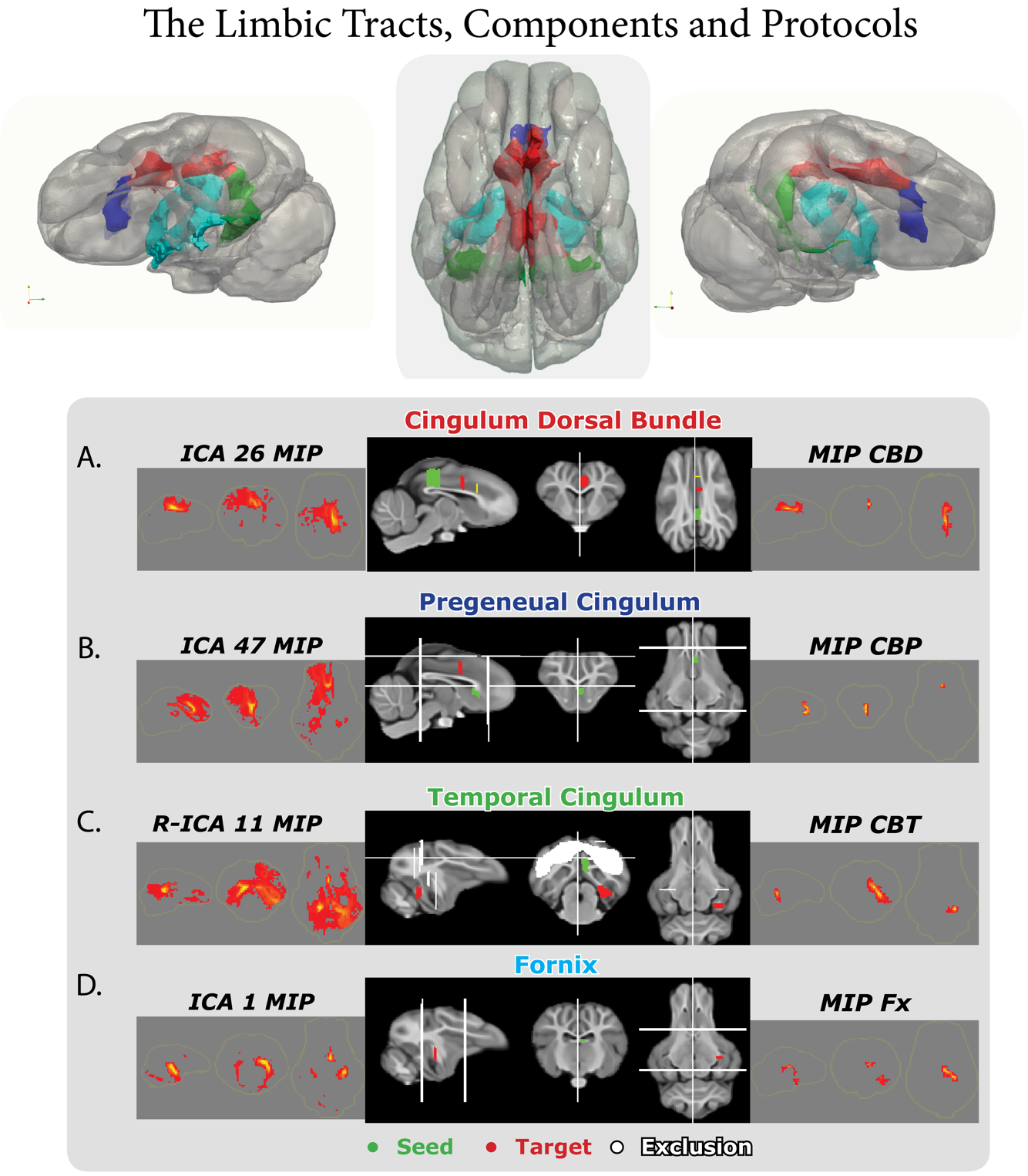

Figure S4: The Limbic tracts of the pig and human, including the Cingulum Dorsal

Bundle (CBD), The Pregenual Cingulum (CBP), the Temporal Cingulum (CBT), and the

Fornix (Fx) reconstructed in 3D.

A). The maximum intensity projection (MIP) of component L-26, the tractography protocol

used for reconstruction of the CBD, and the MIP of the CBD reconstructed with

the mask protocols. B). The MIP of component L-47, the tractography protocol, and

the MIP of the CBP. C). The MIP the component R-11, the tractography protocol, and

the MIP of the CBT. D). The MIP of component L-1, the tractography protocol, and the

MIP of the Fx.

*Cingulum Dorsal Bundle (CBD):* The CBD can be found in components L-26 and R-28 coursing through the cingulate cortex superior to the corpus callosum as the central segment between the temporal and pregenual bundles (Figure 6A). The CBD is seeded in a coronal region of the dorsal posterior cingulate just above the splenium of the corpus callosum. The target mask lies at the border of the anterior and posterior cingulate cortex as defined in the Saikali et al. atlas, and the stop mask is placed just anterior to the target mask (Figure 6A). The tract passes cleanly through both seed and target masks, terminating at the border of the pregenual bundle.

*Pregenual Cingulum (CBP):* The CBP protocol is guided by components L-47 and R-44 and is seeded inferior to the genu of the corpus callosum with the CBD stop mask as its target (Figure 6B). The axial exclusion mask has a slice superior to the genu of the corpus callosum and a second axial slice at the level of the genu of the corpus callosum. A hole is left in the second slice of the exclusion mask, allowing streamlines to propagate throughout the Dorsal Anterior Cingulate. The tract enters the Dorsal Anterior Cingulate and passes the genu of the corpus callosum rostrally curving beneath it to terminate in the anterior prefrontal cortex.

*Temporal Cingulum (CBT):* The CBT is present in component R-11 running parallel to the fornix and connects the dorsal posterior cingulate and parahippocampal cortex in concordance with the work of Bech et al. (Figure 6C)(Bech et al., 2020b). The CBT was seeded in the parahippocampal cortex, with a target placed in the dorsal posterior cingulate, posterior to the splenium of the corpus callosum(Bech et al., 2020b). The exclusion mask blocks the superior hippocampus, visual cortex, and the body and genu of the corpus callosum (Figure 6C). The stop mask is divided into two sections: one, anterior the target, and the second, posterior the fornix.

*Fornix (Fx):* The Fornix is included in the Saikali et al. atlas and is present in components L-01 and R-36/41 (Figure 6D) (Bech et al., 2020b; Saikali et al., 2010b; Zhong et al., 2016a). The fornix is seeded at its apex as defined in the Saikali et al. atlas. It runs inferior to the corpus callosum through the hippocampus and amygdala, terminating in the parahippocampal area anterior to the CBT. The coronal target mask is in the inferior hippocampus**.** The exclusion mask has two coronal slices: one posterior the caudate and the second posterior the genu of the corpus callosum (Figure 6D).

**A Cortex Wide connectivity fingerprint:**

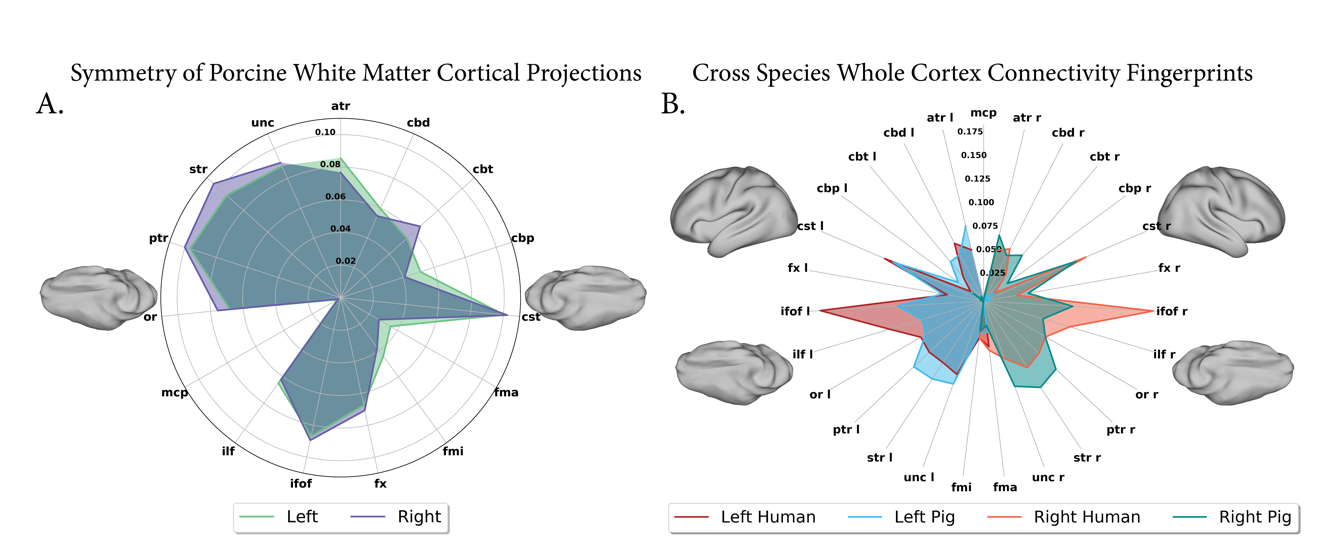

Figure S5) The whole cortex connectivity fingerprints of the pig and human.

A). Symmetry shown via the whole cortex fingerprint of the left and right hemispheres shows the pig to have similar connectivity within each hemisphere. Of note, the prefrontal Forceps Minor (FMI) and Cingulum Pregenual Bundle(CBP) show a bias to the left hemisphere, while the superior and posterior thalamic radiations (STR/PTR) have more robust connectivity in the right hemisphere.

B). Connectivity fingerprints of theleft and right cortex in both species. Connectivity across hemispheres is overall symmetricin both species. The connectivity fingerprint shows which tracts are vital in driving the KL divergence across the pig and human cortex. Of particular note are the increased human connections to the Inferior Fronto-Occipital Fasciculus (IFOF), Inferior Longitudinal Fasciculus (ILF), and Cingulum Dorsal Bundle (CBD). The pig cortex had increased connectivity with the Cingulum Pregenual Bundle (CBP), Uncinate Fasciculus (UNC), and the STR and PTR.
